# Dynamic Behavior of Coupled Neuron System from a Game-Theoretic Standpoint

**DOI:** 10.1101/2020.04.18.046235

**Authors:** Avinash Kori

## Abstract

This paper is concerned with the theoretical investigation of game theory concepts in analyzing the behavior of dynamically coupled oscillators. Here, we claim that the coupling strength in any neuronal oscillators can be modeled as a game. We formulate the game to describe the effect of pure-strategy Nash equilibrium on two neuron systems of Hopf-oscillator and later demonstrate the application of the same assumptions and methods to *N* × *N* neuronal sheet. We also demonstrate the effect of the proposed method on MNIST data to show the equilibrium behavior of neurons in a *N* × *N* neuronal grid for all different digits. A significant outcome of the paper is a modified Hebbian algorithm, which adapts the coupling weights to neural potential resulting in a stable phase difference. Which in turn, makes it possible for an individual neuron to encode input information.

## 1 Introduction

Humans, like many other primitive organisms, are made up of the complex neural system, which is the combination of a large number of neurons forming complex neural architecture. Individual neurons are the building blocks of these complex neural architectures. Many models for single neuron behavior have been proposed and studied, to understand the coupling and firing patterns of these neurons. These models are the set of differential equations derived from the Hodgkin-Huxley model [1]. Some commonly used models are Hopf-field model [2], FitzHugh-Nagumo model [3], Van der pol Oscillator [4], and Morris-Lecar neuron model [5]. These models can capture some behaviors of the biological neurons [6, 7, 8], As described in [9] Morris-Lecar model can capture the behavior of the membrane potential exhibiting spiking, or bursting state by changing the external forced current.

Biological neurons always interact with one another to process information. The coupling in-between the neurons can be a result of electrical or chemical synapses. At equilibrium, these neurons behave in synchrony, which results in a rhythmic group movement of information [10, 11]. This rhythmic movement is the result of neural coupling [10]. Tuning these coupling patterns makes it challenging to analyze the dynamic behavior of the neurons.

Recently, game theory has gained some attention in the field of neuroscience [12]. For instance, Neuroeconomics is the field where the concepts from game theory are extensively used to understand the decision-making process in humans [13]. Game theory can be applied to a wide variety of fields to analyze decision-making strategies followed by players [14]. In [15], the authors formulated a simple two-player strategy game and tried to correlate their model prediction with functional MRI (fMRI) data. Most behavioral game theory literature demonstrates that humans think strategically in a single game. However, it strongly considers the strategies of the opponent [16].

Dynamic game theory has also gained attention in recent days. In a dynamic system (DS) games, the very act of a player changes the dynamics of the game [17]. DS games have an evolving system of differential equations as a game environment, these games have temporal components, and players make decisions at every instant of time. Evolutionary game theory [18] has also offered some similar concepts of using the temporal component in games.

Considering the beliefs as variable and updating the beliefs based on some learning is rarely incorporated in the games. In the context of this paper, we consider every neuron to be a player, and they are provided with the two different strategies which correspond to couple or not to couple with other neurons. The coupling weightage if made as a learnable parameter, which is updated based on the player’s modified Hebbian rule.

Figure 1 describes the overall structure of the proposed framework. In the remainder of this text, Section 2 summarizes the Hopf-Oscillator and outlines circuit dynamics of the coupled neuron system. Section 3 deals with Self-Organizing Maps. Section 4 includes the game-theoretic formulation of the problem, also provides the interpretation of biological neurons. Section 5 summarizes the learning step and outlines its relation with game theory. Section 6 includes experimental details, followed by results and discussion in section 7. Section 8 ends the paper with conclusion.

**Figure 1:**
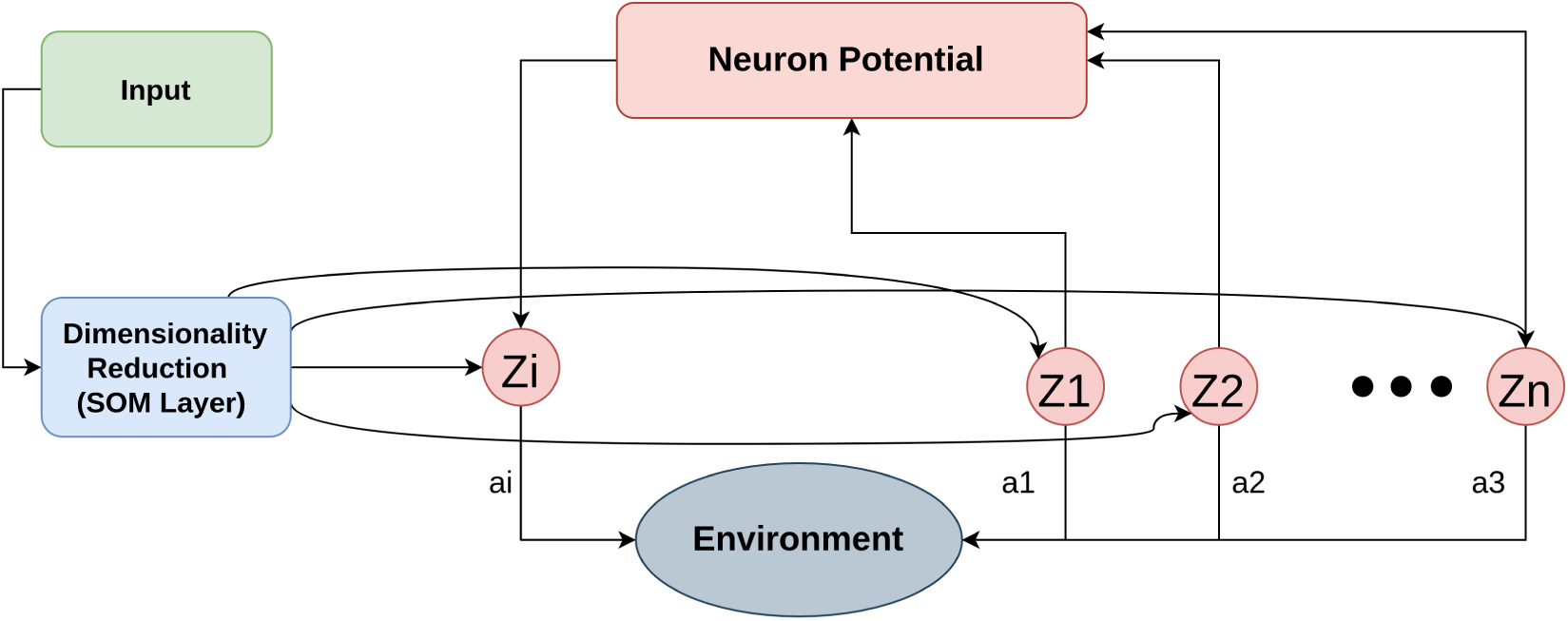
Above figure illustrates the proposed framework. Input is random in the case of two neuron framework, While it’s MNIST digit in case of network example. Dimensionality reduction is the SOM layer for network games while it’s an identity block in the case of two neuron games. *Z*_*i*_ are neurons included in the game, *a*_*i*_ is the action followed by the neuron (player) *i*, the selected action affects the environment, the potential of the neuron affects the firing of other neurons.

## 2 Hopf Coupled Oscillator

As discussed previously, we’ll be using Hopf oscillator for this work. Hopf oscillator is simplified Hodgkin-Huxley model, with just two parameters *µ* and *ω*. Equation 1 shows the single neuron Hopf oscillator, where *z* ∈ ℂ, 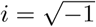, and 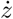 indicates time derivative of complex variable *z*. This simple looking equation exhibits bifurcation property. Figure 2 describes the convergence, divergence, stable state conditions of Hopf oscillator.

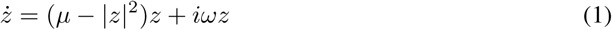

**Figure 2:**
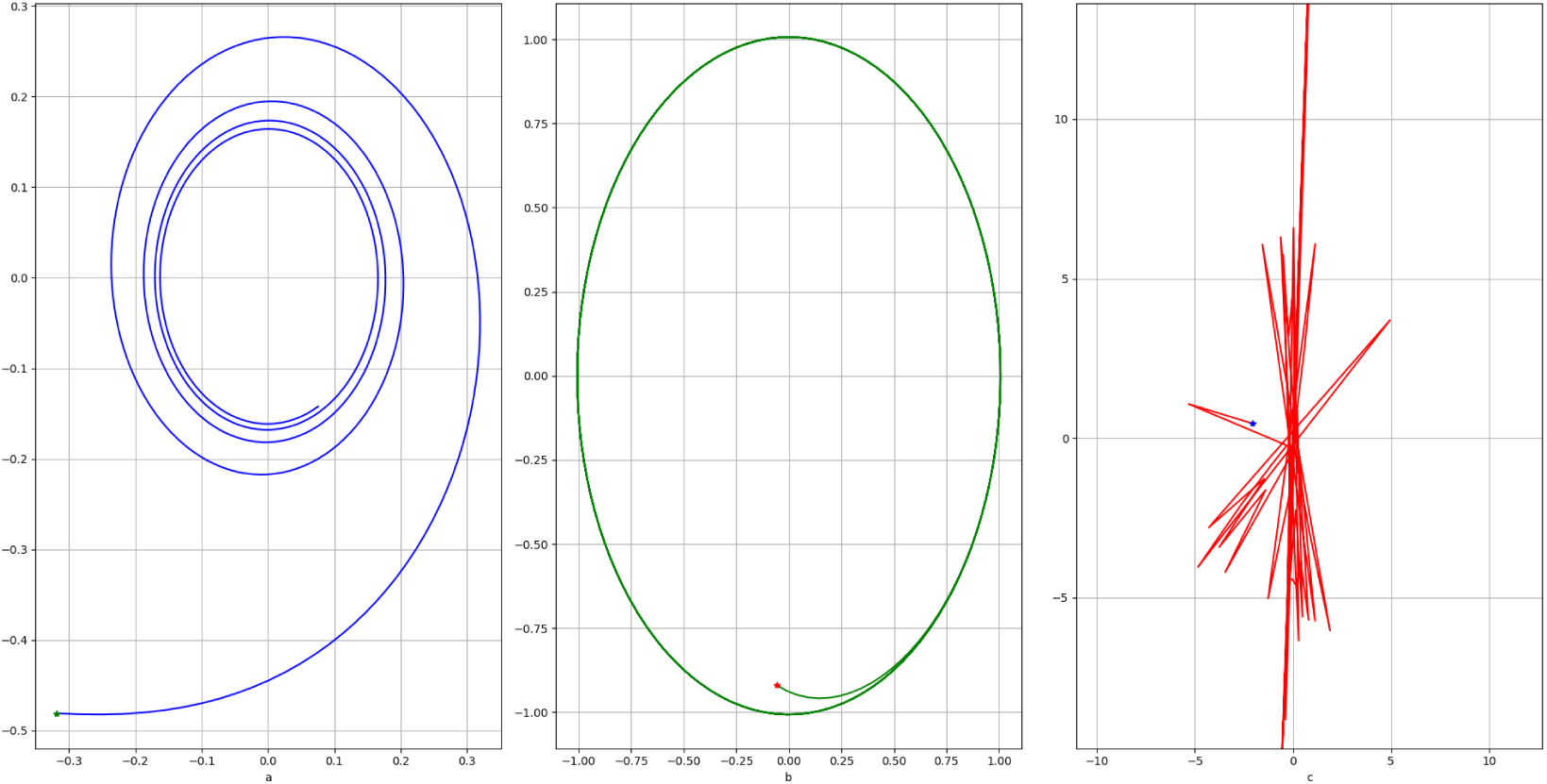
(a) Indicates the convergance of the system (*µ* = 0.1), (b) Indicates the stable state of the system (*µ* = 1.0), and (c) Indicates divergance criteria of the system (*µ* = 15). In all the expirements above 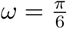, *dt* = 0.1, and * indicates the randomly initialized starting point.

In this work, for the formulation, we make use of a coupled neuron system with two neurons. Figure 3 describes the coupling between two neurons. Let *z*_1_, *z*_2_ be two different neurons; the activity of one is to get coupled with another neuron. Equation 2 describes the coupled neuron system, where *W*_1_, *W*_2_ ∈ ℂ captures interdependent coupling factor.

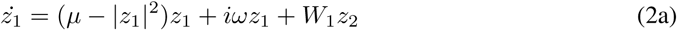

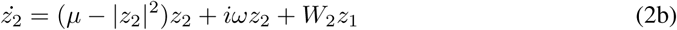

**Figure 3:**
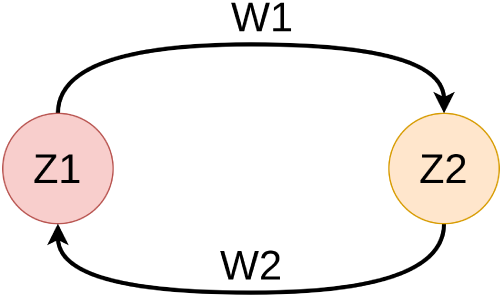
*z*_1_ and *z*_2_ indicates two different neurons, *W*_1_, *W*_2_ indicates the coupling factors for *z*_2_ and *z*_1_

**Figure 4:**
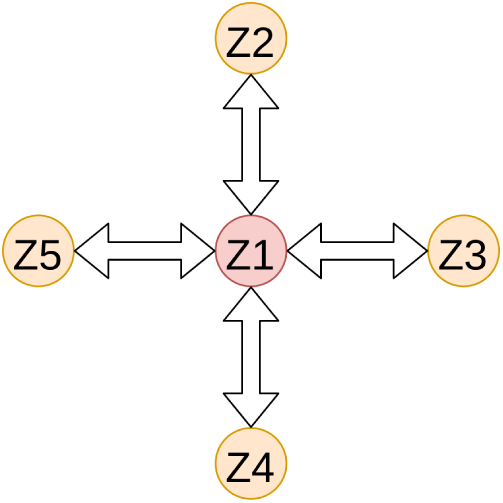
{*z*_*i*_} indicates 5 different neabouring neurons in a network, there exist 2 couple factor inbetween every two neurons

By considering, eular form representation of complex numbers (i.e *z* = *re*^*iθ*^ and *W* = *Ae*^*iϕ*^) and seperating real and imaginary components, the imaginary part of above equations 2 reduces to equation 3 and real part reduces to equation 4

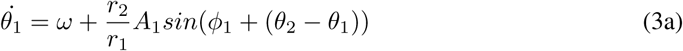

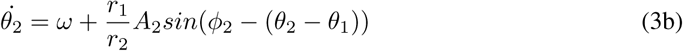

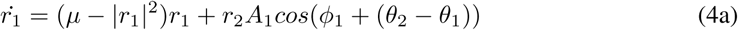

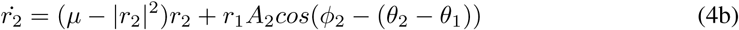

Let (*ψ* = *θ*_2_ − *θ*_1_, *r* = *r*_2_ − *r*_1_), at equilibrium 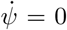,and 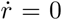 due to syncronization property followed by biological neurons as described in [10, 11].

## 3 Self Organizing Map (SOM)

The proposed network game uses the Self-organizing Map (SOM) model as proposed by Kohonen [19], which accounts for several key features of MNIST handwritten digits [20], these features are further used by the neural sheet for further steps. SOM is a type of unsupervised learning framework used for dimensionality reduction. In our case, we use SOM to reduce the dimensionality of MNIST data from (28 × 28) to 10 × 10. The SOM was trained with MNIST handwritten digits. Once trained, the responses of the SOM layer for any handwritten digits is described in figure 5, we make use of these responses with key features for our further analysis.

**Figure 5:**
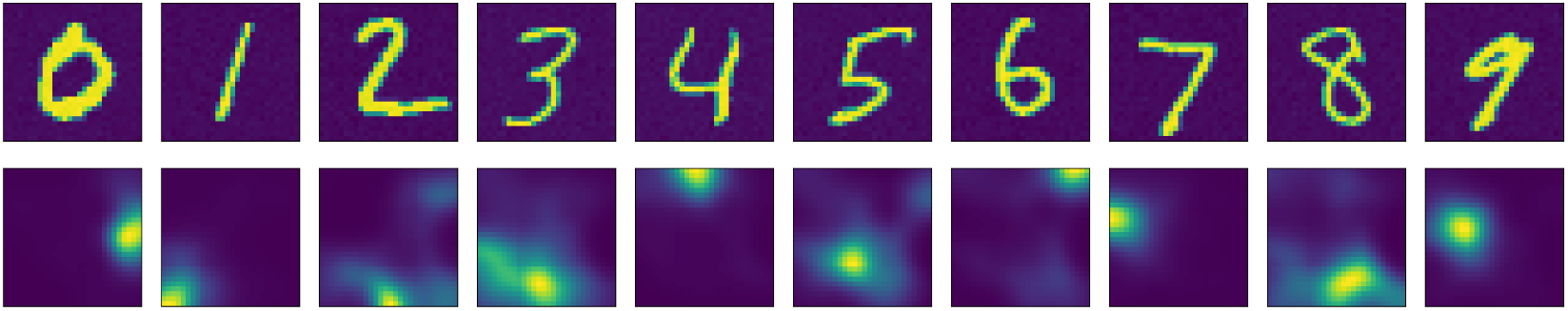
SOM response on MNIST handwritten digits; top row indicates MNIST test digits and bottom row indicates SOM responses for the given input data

## 4 Game Theoretic Formulation and Interpretations

In order to appreciate and fully understand forthcoming sections, it’s important to brief about different games and also layout the proposed structure mathematically. For this work, we formulate this coupling game as a strategic form game. In this game, we assume that the decisions of both the players are independent and non-cooperative in nature. Figure 3, indicates the both players involved: *N* = {*Neuron* − 1, *Neuron* −2}, strategies possible for both the players are: *s*_*i*_ = {*Couple*(*C*), *Don*^*i*^*t* − *Couple*(*DC*)}. The neuronal potential to mapped to strategies as described in equation 5. Strategy set *s*_1_ and *s*_2_, will result in total *s*_1_ × *s*_2_ strategies. In this case all the strategies are: {{*C, C*}, {*C, DC*}, {*DC, C*}, {*DC, DC*}}. As biology encourages synchrony, we model the utilities accordingly; the payoff matrix in figure 6 describes the same. In the payoff matrix in figure 6, using the definitions of rationality and common knowledge, the game can be reduced to an equilibrium point where both the players find coupling as the most advantageous action (strategy set {*C, C*}).

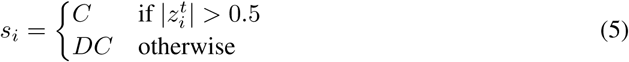

**Figure 6:**
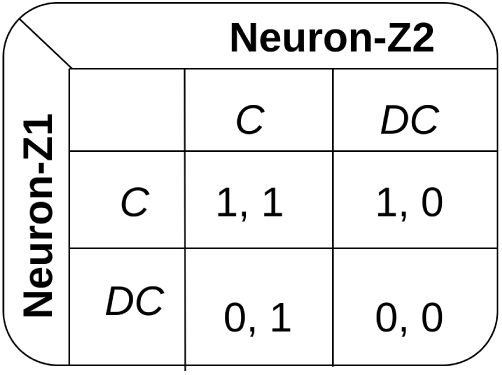
C

### Payoff Matrix

Payoff matrix is used to map set of strategies to there utilities, where utility function is a mapping from strategy space to real-valued number (*u*_*i*_ : {*s*_1_ × *s*_2_} → ℝ). In this paper, we model coupling as a game, figure 6 describes the payoff matrix for this game. As synchronization can be observed from biological neurons, we provide a higher payoff for the player willing to a couple.

### Common Knowledge

Common Knowledge is information that is the same and known by all the players. In the formulated game as any player at any point of time know the utilities and all the strategies of other players, which makes it complete information game. (Every player knows all the possible strategies of other players, but not the exact strategy what the opponents play)

### Rationality

Economists and mathematicians invoke arguments about “rational” behavior to explain the behavior of players based on given circumstances [21]. These assumptions can be considered as building blocks in game theory. One of the assumptions of rationality is that the player never plays a strategy, which provides less payoff when compared to other strategies in certain external circumstances.

### Equilibrium

Is a point in the game where every player selects a strategy and fix to that strategy irrespective of other players. At this state, no player finds it advantages to switch strategy if the strategies of the other players remain unchanged. The set of strategies at this state is considered as equilibrium strategies, More commonly referred to as Nash Equilibrium [22].

### Rationality and equilibrium in the above game

Based on the rationality assumption discussed above, both the players find it optimal to follow the coupling strategy, which results in Pure Strategy Nash Equilibrium at strategy set {*C, C*}. Again by rationality assumptions, it can be easily seen that the strategy set {*C, C*} also survives Iterative Elimination of Strictly Dominated Strategies (IESDC) [23].

### How does a player’s action affect the environment?

As the considered game is dynamic in nature, which means the game itself may converge, diverge, or remain in a stable state based on a player’s actions. Because of this changing environment, to ensure constant coupling, we can have a dynamic payoff matrix or learnable parameter associate with coupling terms, which adjusts the coupling parameter and maintain the environment in the desired state.

## 5 Learning

As discussed in the above section, we have an additional learnable parameter associated with the coupling variable to maintain the environment in the desired state. The most commonly used learning rule for neurons is Hebbian learning [24]. Hebbian learning is a neuroscientific theory which is inspired by the biological neural synaptic adjustment mechanism. Hebbian learning tries to explain both functional and structural plasticity in biological neurons [25]. Here, in this work, we make use of the modified Hebbian method, which considers utilities of the neuron along with neuron state in update rule, as described in equation 6. Where 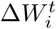 is the weight update for player *i*s coupling variable at that instant *t, u*_*i*_ denotes the utility of player *i, W*^*t*^ indicates the coupling variable at instant *t, z*^*t*^ is neuron potential at instant *t*, and *z*^*t*\*^ is conjugate neuron potential.

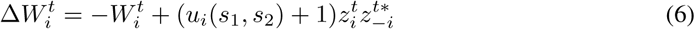

### Equilibrium theoretical analysis

Once the system achieves equilibrium, 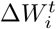 will remain 0, which results in 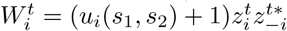.

Converting to Euler form will result in: 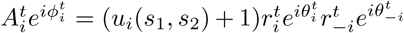, using update terms for both the neurons simultaniously we get equation 7

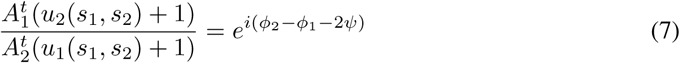

By comparing both the sides we can say that *ϕ*_2_ − *ϕ*_1_ − 2*ψ* = *nπ*, since there’s no imaginary component on left side of the above equation. This phase difference makes the ratio of amplitude of coupling weights propotional to the ratio of utilities as described in equation 8.

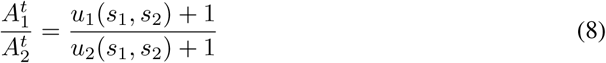

Theoretical coupling to strategies relation is described in the table 1

**Table 1:** Coupling strength based on player strategies

| $s_1$ | $s_2$ | $\frac{A_1^t}{A_2^t}$ |
| --- | --- | --- |
| $C$ | $C$ | 1 |
| $C$ | $DC$ | 2 |
| $DC$ | $C$ | 0.5 |
| $DC$ | $DC$ | 1 |

## 6 Expirements

### Algorithm 1 Proposed method for 2 neuron framework

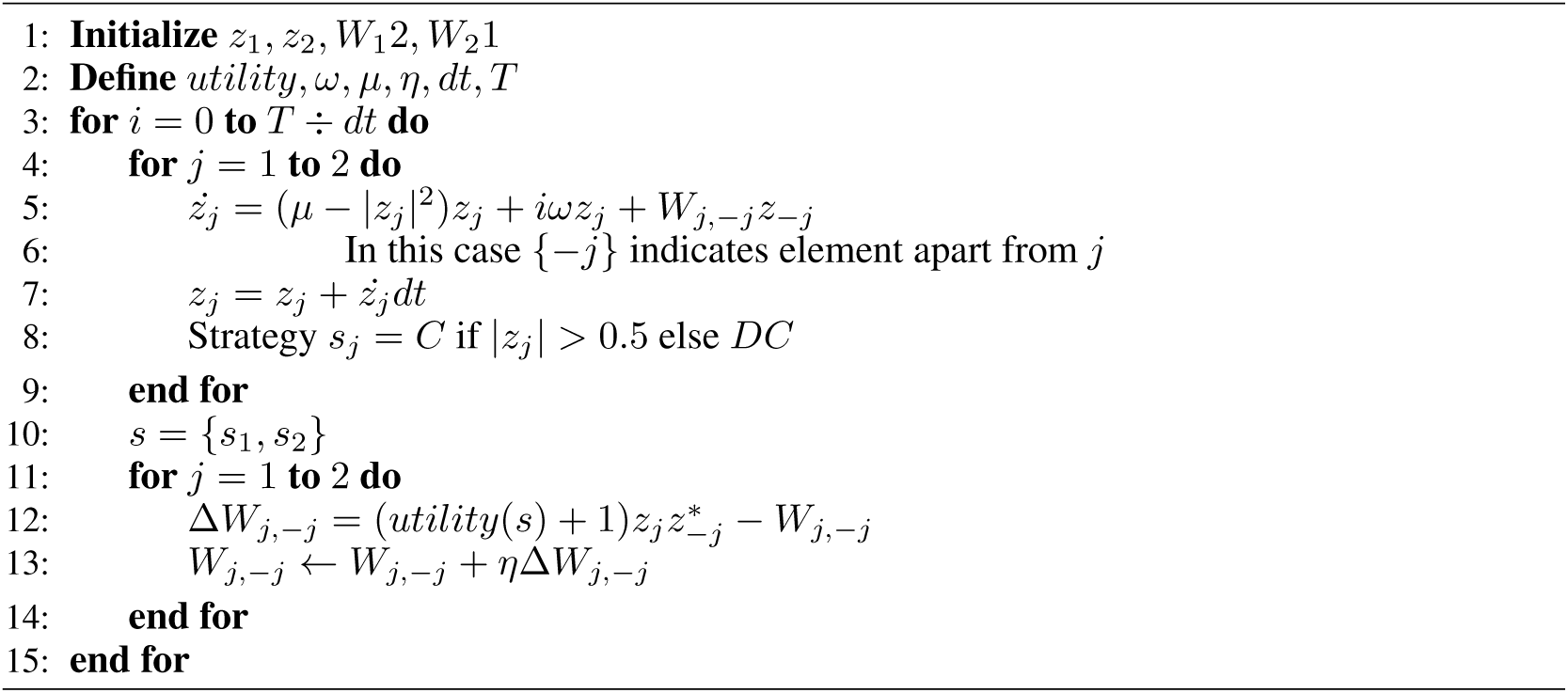

### Two Neuron model

First, to validate the formulated game, we demonstrate the behavior of the neuronal system as described in figure 3, due to coupling and the above utility form. Here, we randomly initialize the potential for *z*_1_ and *z*_2_ and use numerical methods and modified Hebbian rule to drive neuronal potential to an equilibrium point. Algorithm 1 breifly describes the proposed learning method and dynamical system.

### Network Extention

We perform this experiment, to demonstrate the formulated game on the network of *N* × *N* neurons, where each neuron is connected to its adjacent 4 neurons as described in figure 4. We use SOM responses to initialize these neurons and again use numerical methods along with modified Hebbian rule to achieve equilibrium. In this experiment, the aim is to observe different equilibrium coupling phases for different digits in the dataset. The system of equations defining the network game and the corresponding Hebbian rule is described in equations 9 where 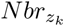 denotes neighboring neurons, which are directly connected to *z*_*k*_.

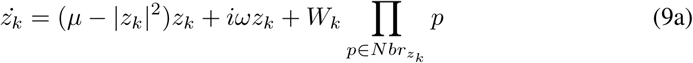

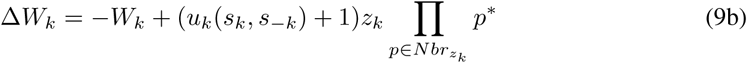

## 7 Results and Discussion

In this study, we show that concepts from game theory can be incorporated in networks to analyze neuronal coupling. As discussed in previous sections, we conducted experiments on two neuron models and network of neurons. In the case of two neurons, it can be seen that the system converges with a fixed phase difference, which is dependent on the random initialization of an oscillator. Figure 7 describes the system oscillation for two different random starting points. In each plot first row illustrates the real part, the second row illustrates the imaginary part, and the third row indicates the phase of the oscillator. In both the figures, it can be observed that both the neurons coupled with the fixed phase difference, which is dependent on the initialization of neurons.

**Figure 7:**
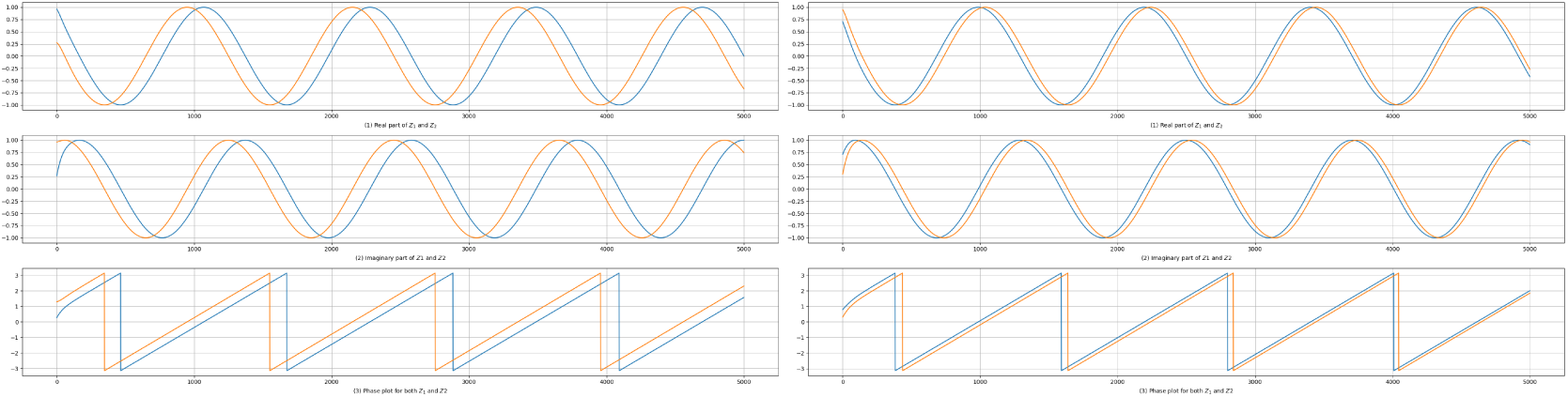
Results for two neuron model, Orange plot denotes player1’s oscillations and blue describes player2’s oscillations

It’s interesting to see that similar behavior can be observed in network oscillations. Figure 8 indicates the unwrapped phase of the player (5,5) in a neural sheet, where it can be easily noted that the oscillator behaves differently for each digit. Similar observations can be done on any randomly selected neuron from a neural sheet 9. This shows that once when the neural sheet oscillates at synchrony, each and every neuron has the capability to differentiate the input, which is very interesting.

**Figure 8:**
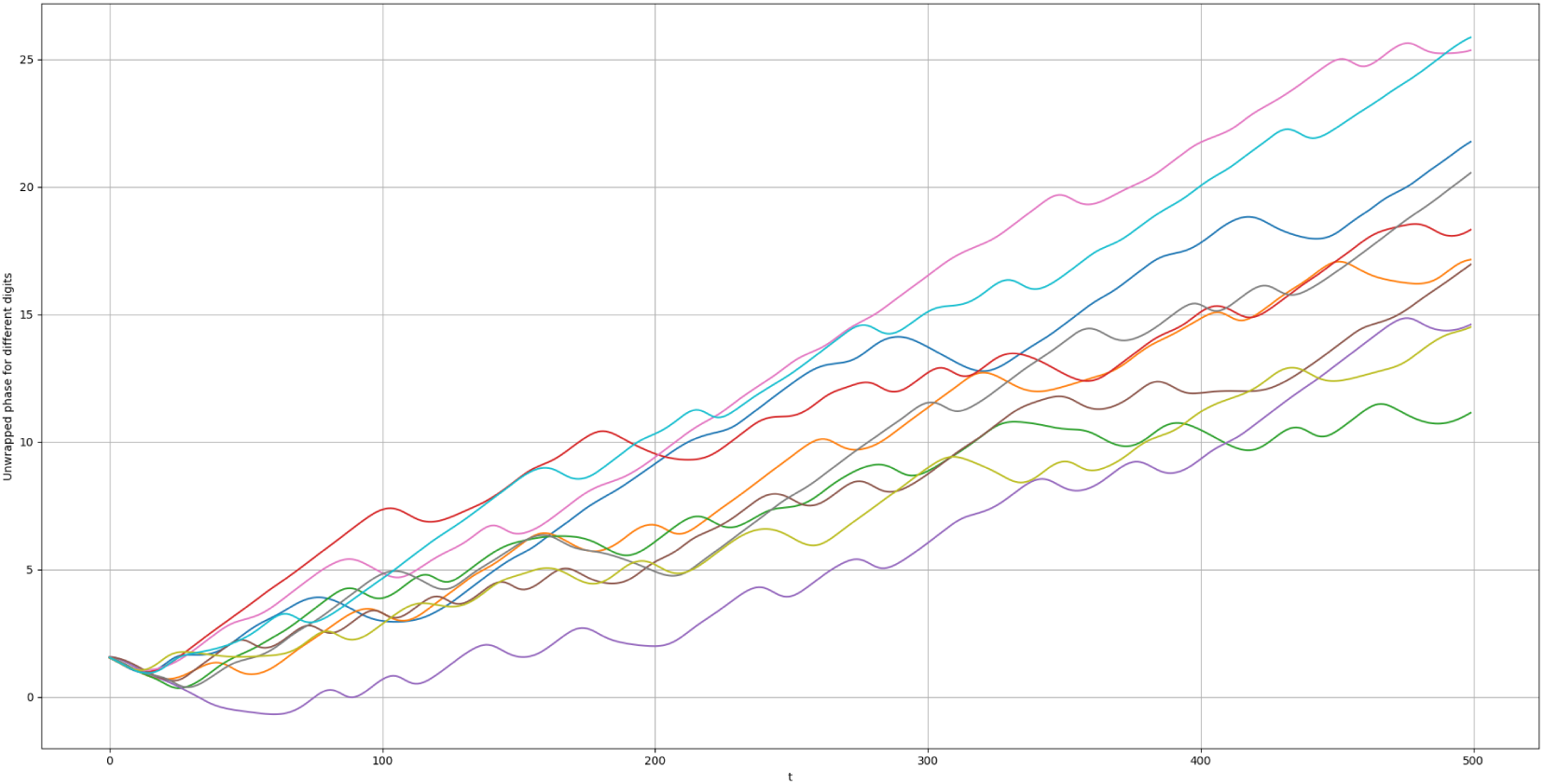
Unwrapped phase of (5,5) neuron in a neural sheet for different input MNIST digit

**Figure 9:**
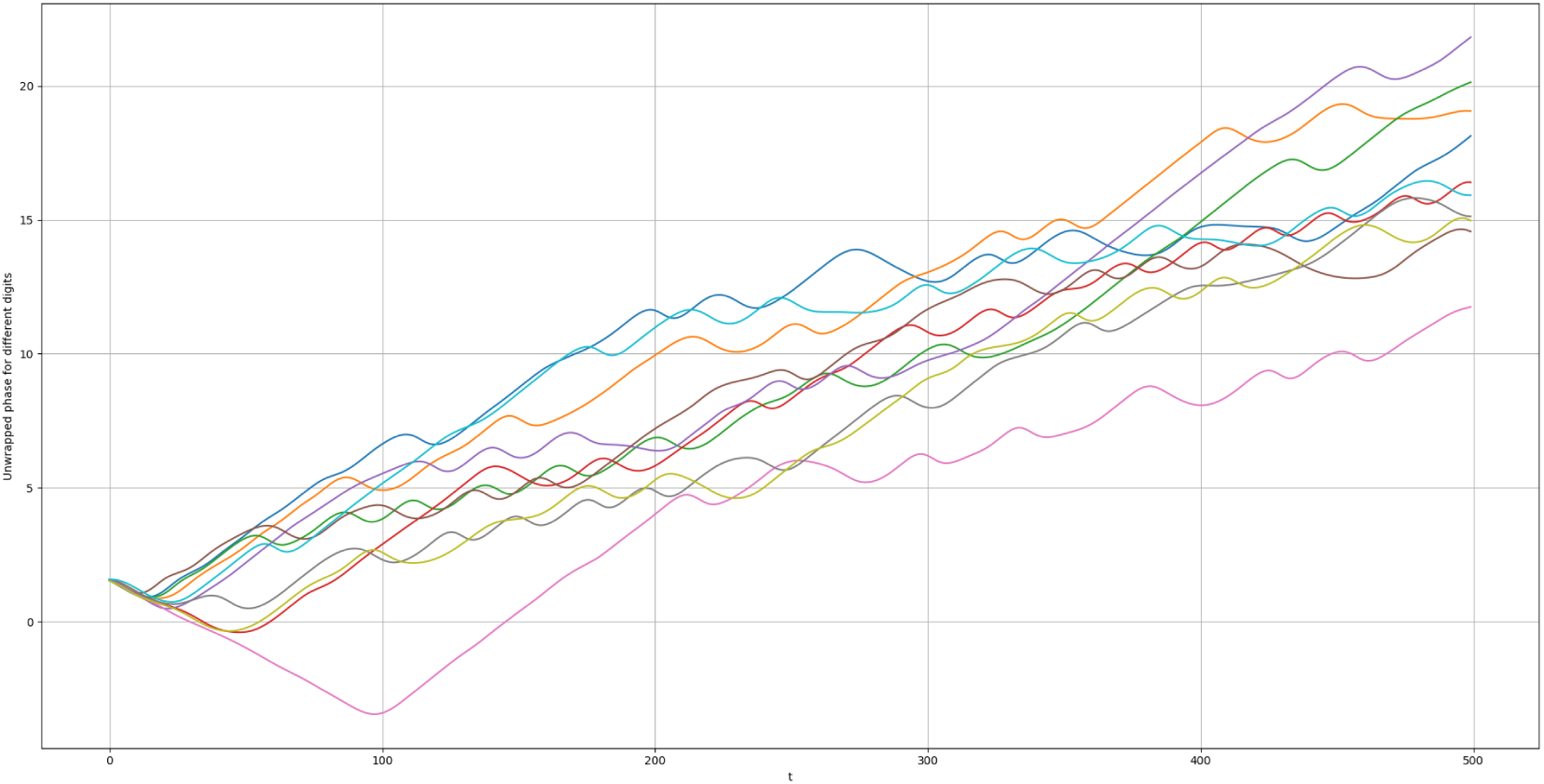
Unwrapped phase of randomly selected neuron in a neural sheet for different input MNIST digit

## 8 Conclusion

The paper provides a novel concept for formulating the neuron coupling as a game, to our knowledge, this is the first time neural coupling is formulated using a game-theoretic framework. In the paper, we also demonstrate the modified Hebbian rule for learning coupling parameters and demonstrate the results on a network and two neuron frameworks. The proposed method is tested on Hopf oscillator but can be easily extended to any other oscillators. These findings are further correlated with biological neuronal behavior in terms of synchronization and synaptic strength development.

## Acknowledgments

The author gratefully acknowledges the discussions with Dr. Srinivas Chakravarthi on neural circuitry, and Dr. Puduru Viswanandha Reddy on the concepts of game theory.

## Notes

### Competing Interest Statement

The authors have declared no competing interest.

